# PMBB Geno-Pheno Toolkit: A suite of scalable, reproducible pipelines for cross-biobank association analyses

**DOI:** 10.64898/2026.07.22.740077

**Authors:** Zachary B. Rodriguez, Lindsay Guare, Lannawill Caruth, Katie. M. Cardone, Christopher Carson, Tess Cherlin, Stephanie Mohammed, Hritvik Gupta, Rachit Kumar, Karl Keat, Shefali S. Verma, Anurag Verma

**Affiliations:** Department of Medicine, Division of Translational Medicine and Human Genetics, University of Pennsylvania - Perelman School of Medicine, Philadelphia, PA 19104, USA; Institute for Biomedical Informatics, University of Pennsylvania - Perelman School of Medicine, Philadelphia, PA 19104, USA; Institute for Translational Medicine and Therapeutics, University of Pennsylvania - Perelman School of Medicine, Philadelphia, PA 19104, USA; Department of Pathology and Laboratory Medicine, University of Pennsylvania, Philadelphia, PA 19104; Jacobs School of Medicine and Biomedical Sciences at the University at Buffalo, Buffalo, NY 14203; Department of Genetics, University of Pennsylvania - Perelman School of Medicine, Philadelphia, PA 19104, USA; Division of Informatics, Department of Biostatistics, Epidemiology, and Informatics, University of Pennsylvania, Philadelphia, PA 19104

**Author notes:** These authors contributed equally to this work (Zachary B. Rodriguez, Lindsay Guare, Lannawill Caruth).

## Abstract

**Summary:** Electronic health record (EHR)-linked biobanks generate unprecedented genomic and phenotypic datasets, but their scientific utility is constrained by data fragmentation across institutional silos and incompatible computing infrastructures, forcing researchers to rewrite ad-hoc scripts for each new environment. We present the PMBB Geno-Pheno Toolkit, a suite of modular Nextflow pipelines for biobank-scale association analyses. This note focuses on the toolkit’s SAIGE family of pipelines — supporting genome-wide (GWAS), exome-wide (ExWAS), and phenome-wide (PheWAS) association testing — together with the companion GWAMA and ExWAS meta-analysis pipelines that enable cross-biobank replication. All components are containerized (Docker/Apptainer) and orchestrated with Nextflow, allowing the same workflows to run unmodified on local HPC clusters, cloud platforms, and the All of Us Research Workbench. Complementary toolkit pipelines for PLINK-based GWAS, polygenic scoring, LD-based clumping, and phenotype harmonization are also available and briefly noted.

**Availability:** The PMBB Geno-Pheno Toolkit is freely available at https://github.com/PMBB-Informatics-and-Genomics/pmbb-geno-pheno-toolkit under MIT open-source license.

## Introduction

The electronic health record (EHR)-linked biobank era has reached a transformative milestone, amassing data from millions of participants across global population-based and academic medical centers, along with large-scale genotyping, exome, and whole-genome sequencing datasets (Kurki *et al*., 2023; Sudlow *et al*., 2015; Verma *et al*., 2022; All of Us Research Program Genomics Investigators, 2024). While these repositories offer unprecedented potential to pinpoint genetic variations and unravel phenotypic complexities, their scientific utility is severely constrained by a pervasive innovation-application gap. Current genomic discovery is hampered by data fragmentation across institutional silos and incompatible computing architectures, which prevents the systematic meta-analysis and cross-population replication required to validate findings and identify low-frequency associations (Verma *et al*., 2024).

This fragmentation is largely driven by a lack of standardized, reproducible, and portable analytical frameworks. Biobank-scale studies currently rely on ad-hoc scripts and custom workflows that must be rewritten for every new computing environment, from local high-performance computing (HPC) clusters to cloud platforms. This reliance on non-transferable code forces researchers to expend critical resources on reimplementing scaling strategies and debugging platform-specific configurations. These technical burdens propagate errors and cast doubt on reproducibility (Leitão, 2004). Standardized workflows for biobank-scale associations and post-association analyses have lagged significantly behind other sequencing fields that have adopted pipeline management systems like Nextflow (Di Tommaso *et al*., 2017) and Snakemake (Köster and Rahmann, 2012), creating a substantial barrier for researchers without extensive bioinformatics training.

To address these shortcomings, we present the PMBB Geno-Pheno Toolkit, a comprehensive suite of modular Nextflow pipelines engineered for scalable, reproducible genomic research across heterogenous biobanks. This application Note focuses on the toolkit’s SAIGE family of association pipelines and the companion GWAMA and ExWAS meta-analysis pipelines, which together support the transition from single-biobank discovery to cross-biobank replication. Complementary pipelines in the toolkit such as Plink-based GWAS, multi-ancestry polygenic risk scoring, LD-based variant clumping, variant annotation, and phenotype harmonization, are also available and noted briefly where relevant but outside the scope of the detailed description below.

## Implementation

The PMBB Geno-Pheno Toolkit is designed to accelerate genomic discovery by transforming fragmented biobank data into actionable scientific insights through a modular, platform-agnostic framework (Figure 1).

**Figure 1.**
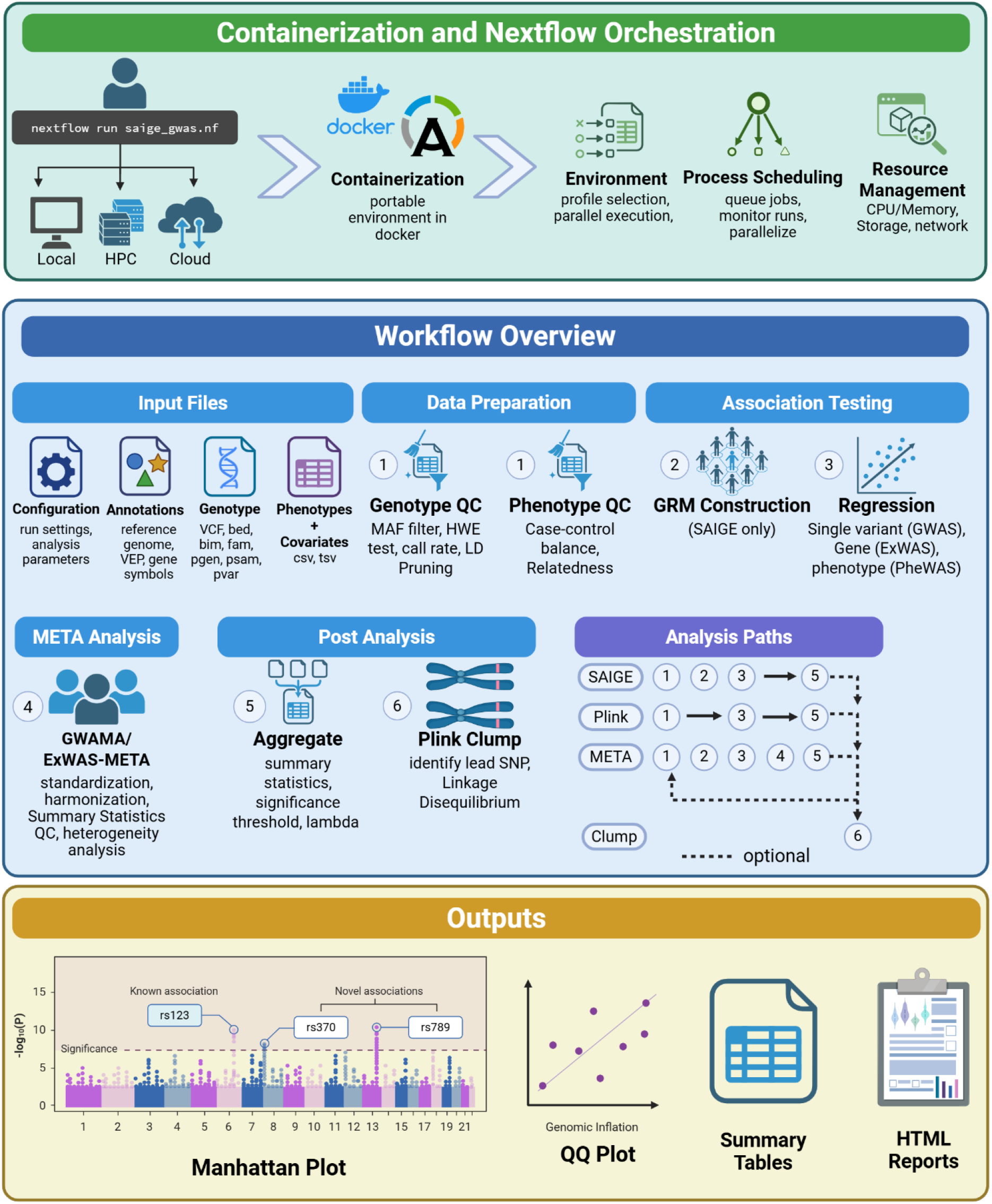
A three-panel schematic of our streamlined bioinformatics pipelines designed to conduct large-scale genetic studies by integrating containerization and automated orchestration. (A) Containerization and Nextflow orchestration: A user can use our pipeline on local workstation, HPC computing environment, and on cloud infrastructure. Our code is available on GitHub and the environment needed to run our pipelines is containerized in Docker and Apptainer. The user chooses a profile to run their analysis and Nextflow handles the parallelization, process scheduling, job scheduling, and resource management. (B) Workflow overview shared by our pipelines: The modular pipelines ingest genomic and phenotypic data specified by the user via simple configuration files, and the steps of the analysis pipelines are executed asynchronously. The pipelines perform rigorous quality control, association testing, and optional meta-analyses and LD-based clumping and pruning. (C) Diagrammatically simplified examples of our pipeline outputs including graphs, plots, summary tables, and HTML reports.

### SAIGE-family association pipelines

The pipelines described in this note perform GWAS, ExWAS, and PheWAS (single-variant and gene-burden) using SAIGE (Zhou *et al*., 2018), a mixed-model method that robustly accounts for population structure and case-control imbalance in biobank-scale data. Within these pipelines, PLINK (Chang *et al*., 2015) is used internally while SAIGE performs the association test and genetic relatedness matrix construction. This is distinct from the toolkit’s separate GWAS with PLINK 2.0 pipeline, which performs association testing directly in PLINK for users who prefer that approach over SAIGE’s mixed model framework.

### Meta-analysis pipelines

SAIGE-family pipelines emit output channels containing summary statistics that feed directly into the GWAMA (Mägi and Morris, 2010) and ExWAS-Meta pipelines, enabling cross-biobank meta-analysis without manual reformatting between steps. A separate post-association pipeline for linkage disequilibrium (LD)-based variant clumping and pruning is also available within the toolkit, allowing investigators to identify independent signals and physically overlapping loci for downstream functional interpretation, but is likewise outside this note’s scope.

### Phenotyping pipeline

A common barrier for researchers new to biobank-scale analysis is that the SAIGE-family pipelines expect phenotype and covariate data as a single wide table (samples × columns); institutional data arrives in mixed formats, spread across multiple source files, with mixed data types and inconsistent encodings across covariates and phenotypes. To lower this barrier, the toolkit additionally provides an optional Phenotyping Pipeline (https://github.com/PMBB-Informatics-and-Genomics/pmbb-nf-toolkit-phenotyping) that aggregates quantitative, categorical, and diagnostic phenotypes from various of source tables and formats into the wide- or long-format table expected by the association pipelines above. Each biobank’s phenotypes and their source tables are described in a single YAML configuration file, which the pipeline resolves into per-phenotype configurations and processes in parallel. It is an optional preprocessing step upstream of the association pipelines described above, is not benchmarked in this note, and is intended to give users without a data-wrangling background a documented path to an analysis-ready phenotype table.

### Reproducibility

To ensure high-fidelity reproducibility across diverse research environments, all analytical components are containerized using Docker and Apptainer. This allows the same scientific workflows to be deployed on local workstations, high-performance computing (HPC) clusters, or cloud-based platforms (e.g., AWS, Azure, Google Cloud) without modification. By leveraging Nextflow’s asynchronous execution, the toolkit removes the performance overhead typical of large-scale genomic studies, enabling downstream post-processing and publication-ready visualizations, including Manhattan plots, QQ plots, and comprehensive association reports to begin as soon as upstream results are emitted.

### Use Case

As a representative use case, we report the performance and portability of the toolkit’s SAIGE GWAS pipeline using Nextflow’s built-in reporting framework to capture resource utilization and runtime across computing environments. Because all toolkit pipelines share the same underlying Nextflow architecture, containerization strategy, and input/output conventions, the runtime and portability characteristics observed here are broadly representative of the SAIGE-family, GWAMA, and ExWAS-meta pipelines described above. Benchmarking datasets included test datasets available on our GitHub and real biobank-scale data from Penn Medicine BioBank (N=56,367 samples; 164,628,091 variants) and *All of Us* (N=10,000 samples; 111,404,689 variants), representative of EHR-linked biobank studies. GWAS using the SAIGE workflow was completed for four phenotypes (two binary: abdominal aortic aneurysm and type II diabetes, and two quantitative: BMI and LDL cholesterol). The analysis on a local HPC cluster completed in 249.7 CPU-hours, with a total runtime of 136 minutes under Nextflow’s asynchronous execution. The parallelization strategy scaled up to 500 concurrent processes as defined in our configuration, with memory usage for all but one task falling below 15GB per worker; one combined plotting task required 56GB. Task execution time ranged from 0–20 minutes (mean: 4.6 minutes for the longest process). On the *All of Us* Research Workbench, we achieved similar results (164 minutes, 357 CPU-hours). Reports for the runs, including summary tables, phenotype distribution plots, Manhattan plots, QQ plots, and analysis logs, are shown in Supplementary Materials. The pipeline successfully processed real-world datasets from 2 institutions, demonstrating robustness across diverse data formats and analytical requirements.

## Conclusion

The PMBB Geno-Pheno Toolkit addresses the systematic fragmentation that continues to hinder large-scale genomic research by providing standardized, portable pipeline infrastructure for high-throughput analysis. While biological data and formats are largely standardized, cross-biobank analysis remains technically impractical due to incompatible computing environment. The toolkit resolves this by shifting the focus from the data itself to the underlying infrastructure, allowing researchers to bypass institutional silos and heterogenous platforms. By wrapping established analytical tools such as SAIGE, PLINK, and GWAMA, within robust Nextflow frameworks, we directly mitigate the challenges highlighted previously: scalability, portability, and reproducibility, reducing the startup costs and technical overhead that stifle genomic research progress. A central goal of the toolkit is to democratize access to sophisticated genomic methods: Our low-code approach lowers the barrier to entry for clinical researchers without extensive bioinformatics expertise to execute rigorous, biobank-scale analyses.

Core association workflows support GWAS, ExWAS, and PheWAS (both single-variant and gene-burden) utilizing SAIGE (Zhou *et al*., 2018) and PLINK (Chang *et al*., 2015) to robustly account for population structure and case-control imbalance in biobank-scale datasets. The association pipelines generate output channels containing summary statistics that serve as direct inputs for the GWAMA (Mägi and Morris, 2010) and ExWAS-Meta (Mägi and Morris, 2010) pipelines, facilitating cross-biobank analyses. Beyond the SAIGE-family and meta-analysis pipelines, the toolkit includes complementary pipelines for PLINK-based GWAS, LD-based clumping, phenotype harmonization, multi-ancestry polygenic risk score calculation (Ruan *et al*., 2022), Variant Effect Predictor (VEP; McLaren *et al*., 2016), giving investigators a consistent path from initial discovery through downstream applications. To support community adoption, we provide comprehensive documentation, example datasets, and tutorial materials through our GitHub repository (https://github.com/PMBB-Informatics-and-Genomics/pmbb-geno-pheno-toolkit).

One of the toolkit’s strengths lies in its modular design, which allows independent analytical components to be combined into comprehensive super-pipelines. Each workflow is built using standard Nextflow processes that utilize integrated data channels (take and emits) to facilitate seamless transfer of information between distinct analytical stages (Di Tommaso *et al*., 2017). This interoperability was engineered to support large-scale association studies, locus characterization, and meta-analyses across cohorts and biobanks. Beyond technical efficiency, the toolkit promotes transparent science, fosters collaboration, and enables independent validation through containerized platform-agnostic pipelines, bringing us one step closer to a decentralized federated model for robust, fair, and sustainable scientific data analysis (Sharma *et al*., 2025).

Several operational constraints remain that we are actively addressing. The implementation does not yet fully conform to nf-core community standards, which may temporarily impact interoperability within the broader Nextflow ecosystem; work towards full compliance is underway. Computational scalability remains a significant hurdle for exceptionally large studies, particularly analyses involving multi-million sample datasets. Data harmonization, specifically non-standardized column naming conventions across disparate institutions, may require manual intervention despite regular-expression matching strategies implemented to catch these issues early in each run. Deployment within certain research environments, such as the UK Biobank DNAnexus Research Analysis Platform, currently requires additional platform-specific configuration. To expand the toolkit’s analytical breadth, we are developing additional modules for hene-based transcriptome-wide association testing with MultiXcan (Barbeira et al., 2019), mendelian randomization, fine-mapping (Gao and Zhou, 2024), LDSC genetic correlation and heritability estimation (Nam et al., 2023), and OMOP common-data-model-based phenotyping.

