## Supplementary Materials for "PMBB Geno-Pheno Toolkit: A suite of scalable, reproducible pipelines for cross-biobank association analyses"

### SAIGE GWAS Results Report

This report presents findings from a SAIGE GWAS (Genome-Wide Association Study) analysis. Single-variant association tests were performed across the specified phenotypes and cohorts.

#### Getting Started

- Use the **Results Filter** in the sidebar to navigate to a phenotype and cohort result.
- Check the **Phenotype Summary** for trait distributions and sample counts.
- Each result page shows Manhattan and QQ plots alongside top hits above the suggestive threshold.
- Refer to **Analysis Logs** for the full pipeline parameter set.

### Phenotype Summary

#### Phenotype Summary Statistics

| COHORT | PHENO | N | Controls | Cases | Prevalence |
|---|---|---|---|---|---|
| PMBB_AFR_ALL | AAA | 12042 | 11744.0 | 298.0 | 0.024746719 |
| PMBB_AFR_ALL | T2D | 12042 | 7093.0 | 4949.0 | 0.410978242 |
| PMBB_AFR_ALL | bmi_median | 11744 | nan | nan | nan |
| PMBB_AFR_ALL | chol_ldl_median | 9083 | nan | nan | nan |
| PMBB_EUR_ALL | AAA | 41651 | 40128.0 | 1523.0 | 0.036565748 |
| PMBB_EUR_ALL | T2D | 41651 | 32886.0 | 8765.0 | 0.210439125 |
| PMBB_EUR_ALL | bmi_median | 38555 | nan | nan | nan |
| PMBB_EUR_ALL | chol_ldl_median | 23164 | nan | nan | nan |

### Phenotype Distribution Plots

## T2D

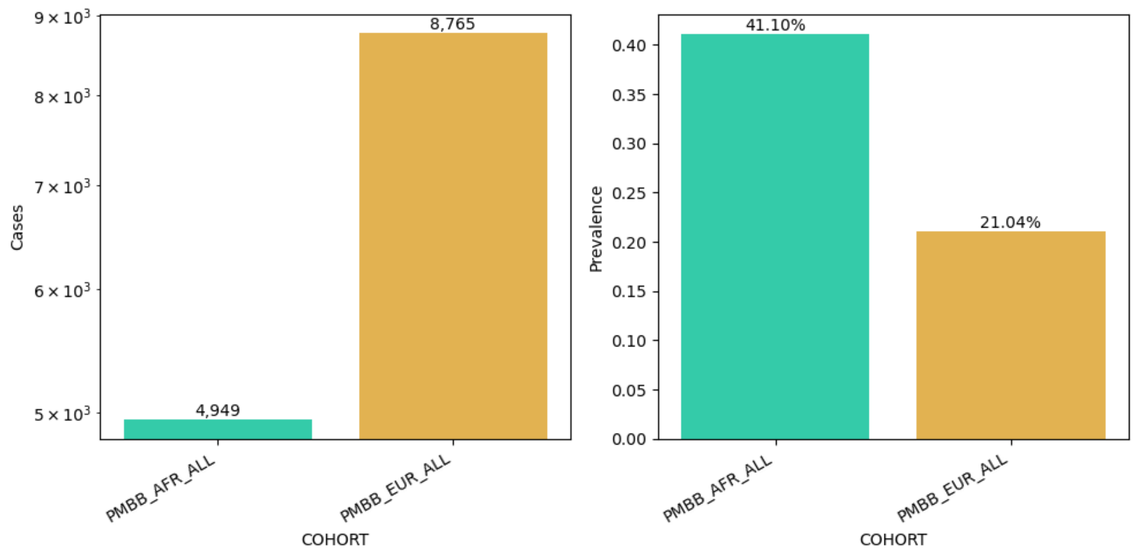

### AAA

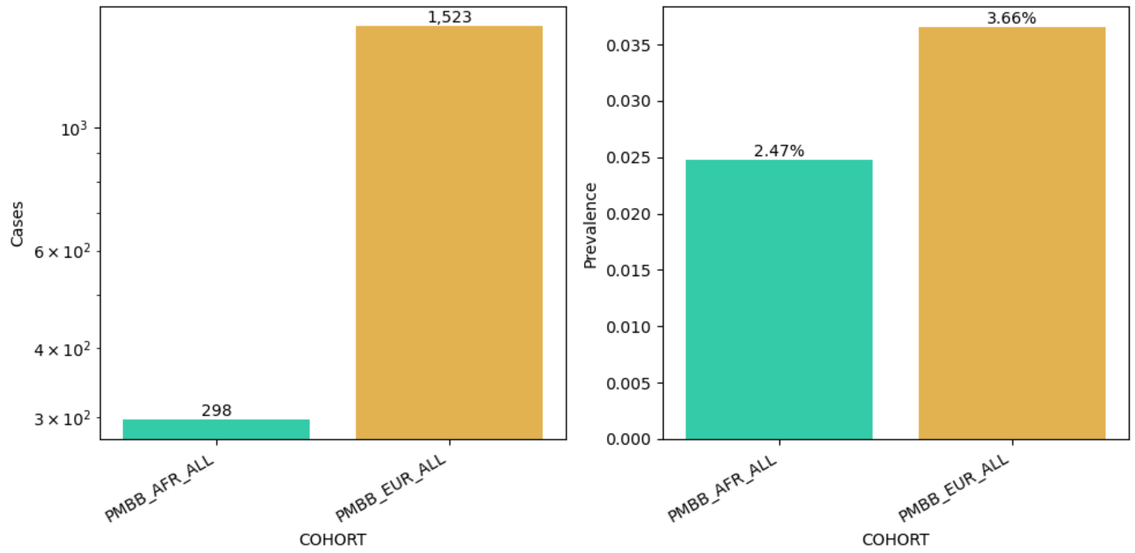
1,523
298
$10^3$
$6 \times 10^2$
$4 \times 10^2$
$3 \times 10^2$
Cases
PMBB_AFR_ALL
PMBB_EUR_ALL
COHORT
3.66%
2.47%
0.035
0.030
0.025
0.020
0.015
0.010
0.005
0.000
Prevalence
PMBB_AFR_ALL
PMBB_EUR_ALL
COHORT

#### chol_ldl_median

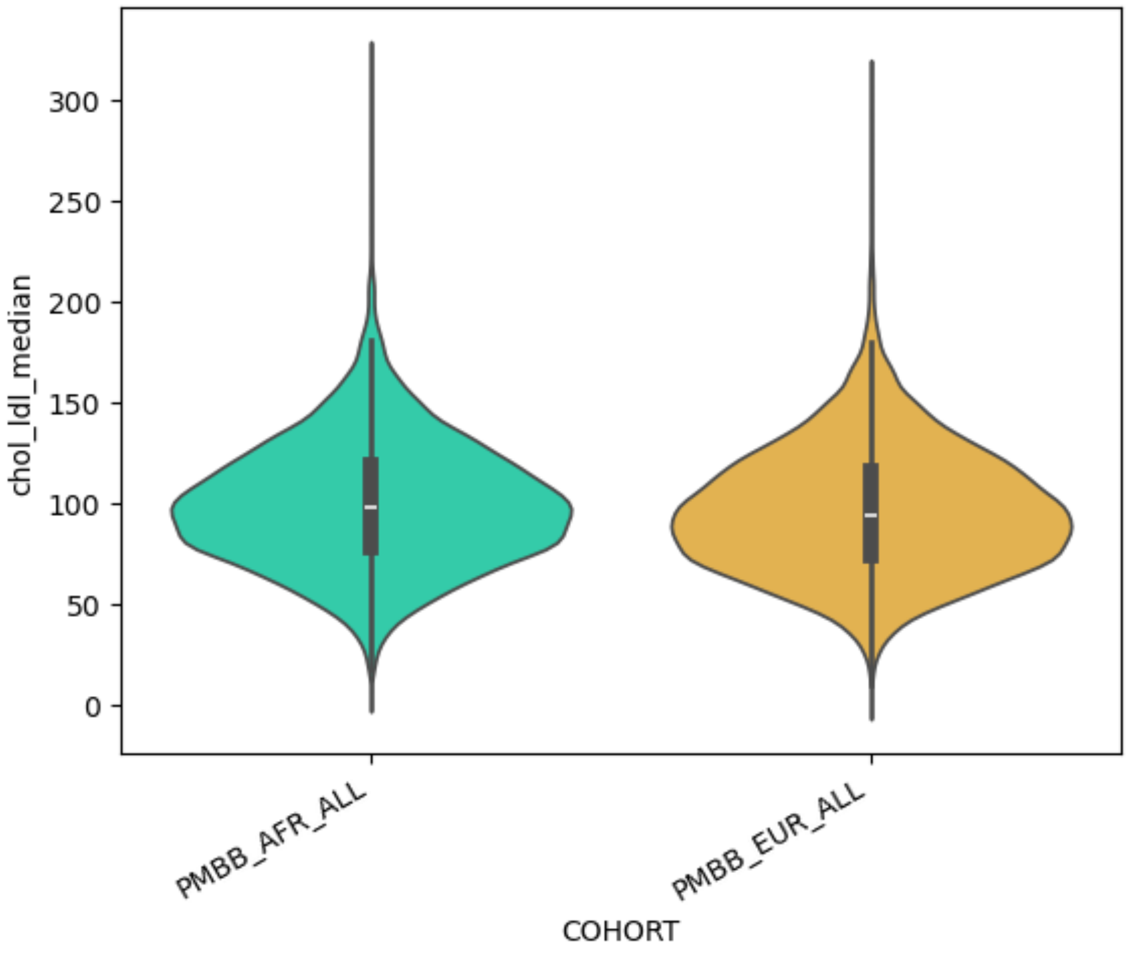

#### bmi_median

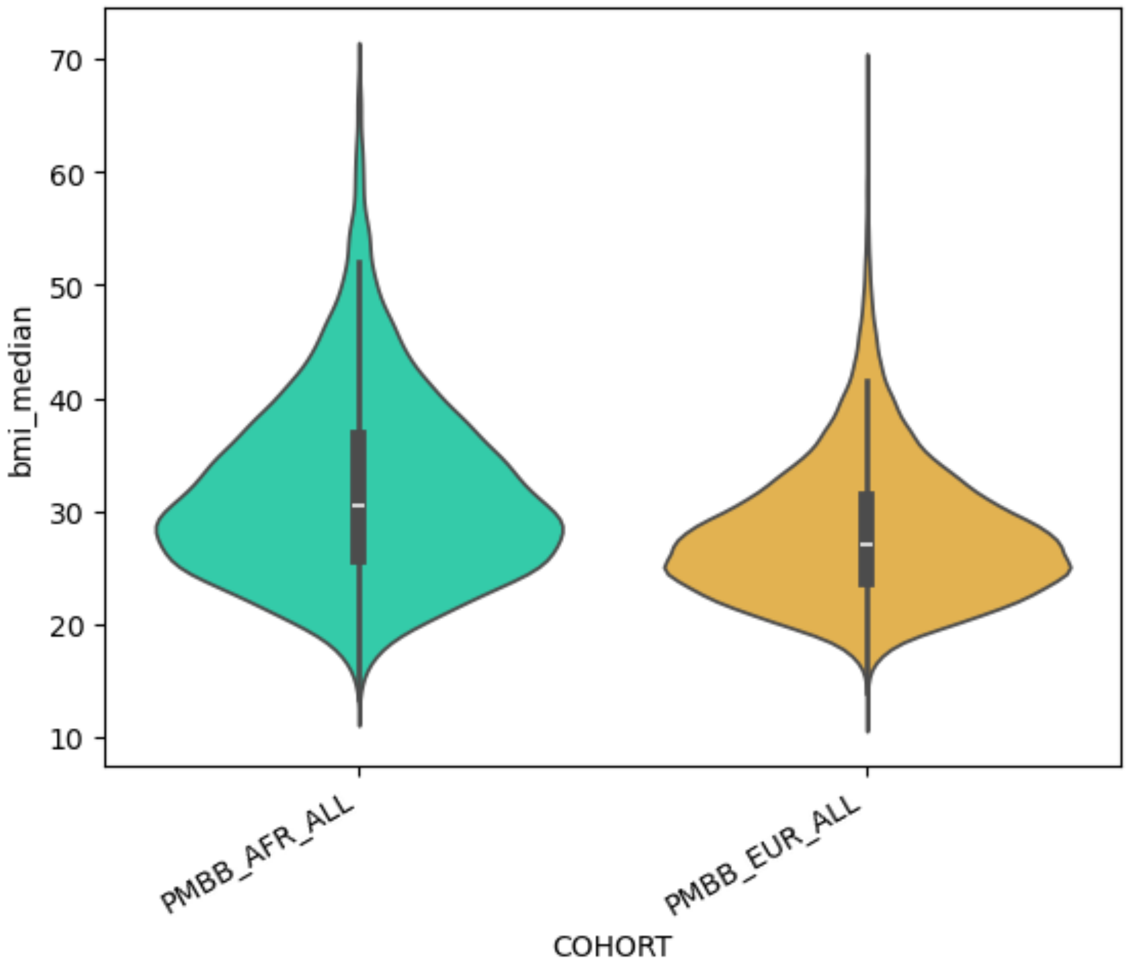

### GWAS — T2D in PMBB_AFR_ALL

#### GWAS Results — T2D in PMBB_AFR_ALL

##### GWAS Plots — T2D in PMBB_AFR_ALL

Manhattan Plot

QQ Plot

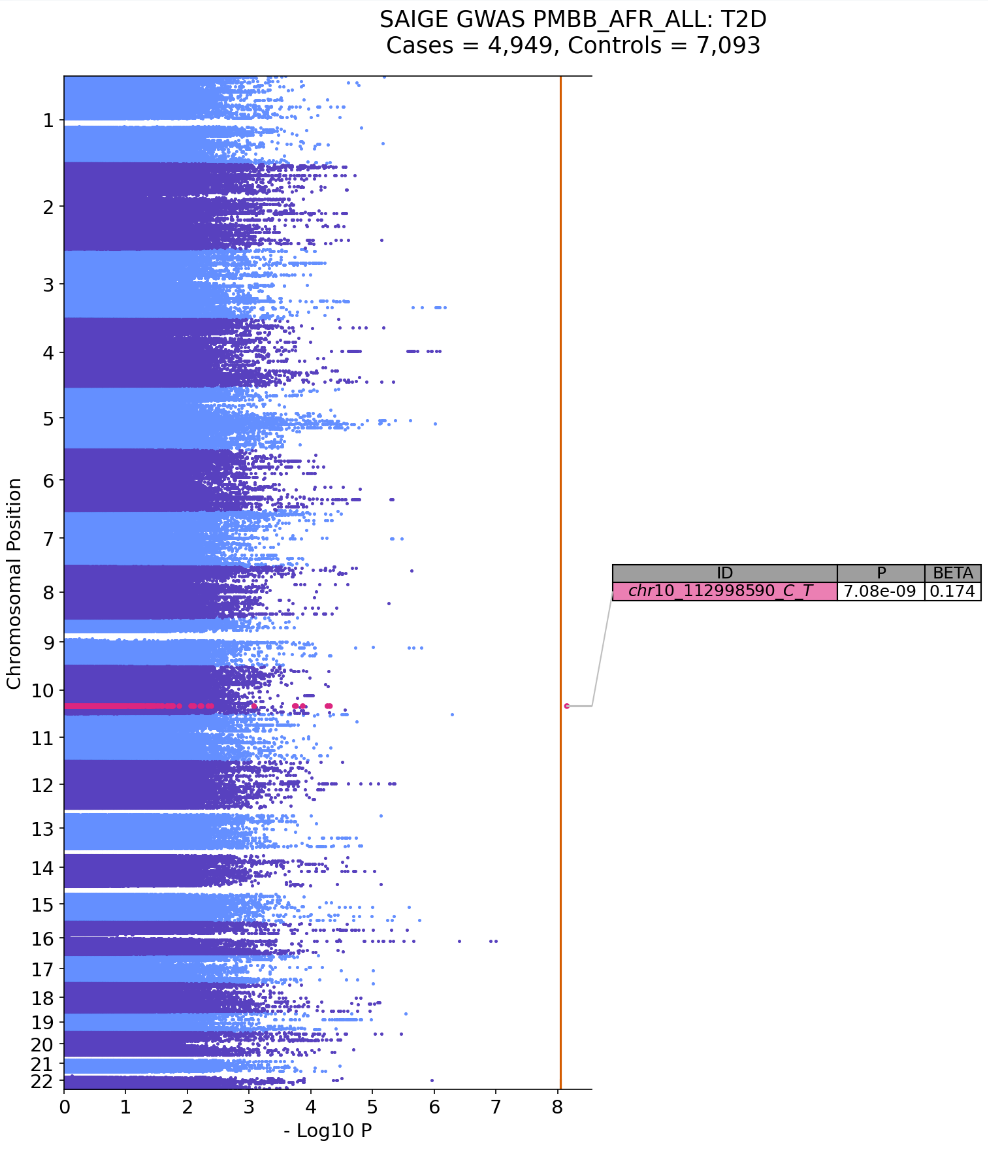


### Top GWAS Hits

No significant hits found above the p-value threshold.

### GWAS — T2D in PMBB_EUR_ALL

#### GWAS Results — T2D in PMBB_EUR_ALL

##### GWAS Plots — T2D in PMBB_EUR_ALL

Manhattan Plot

QQ Plot

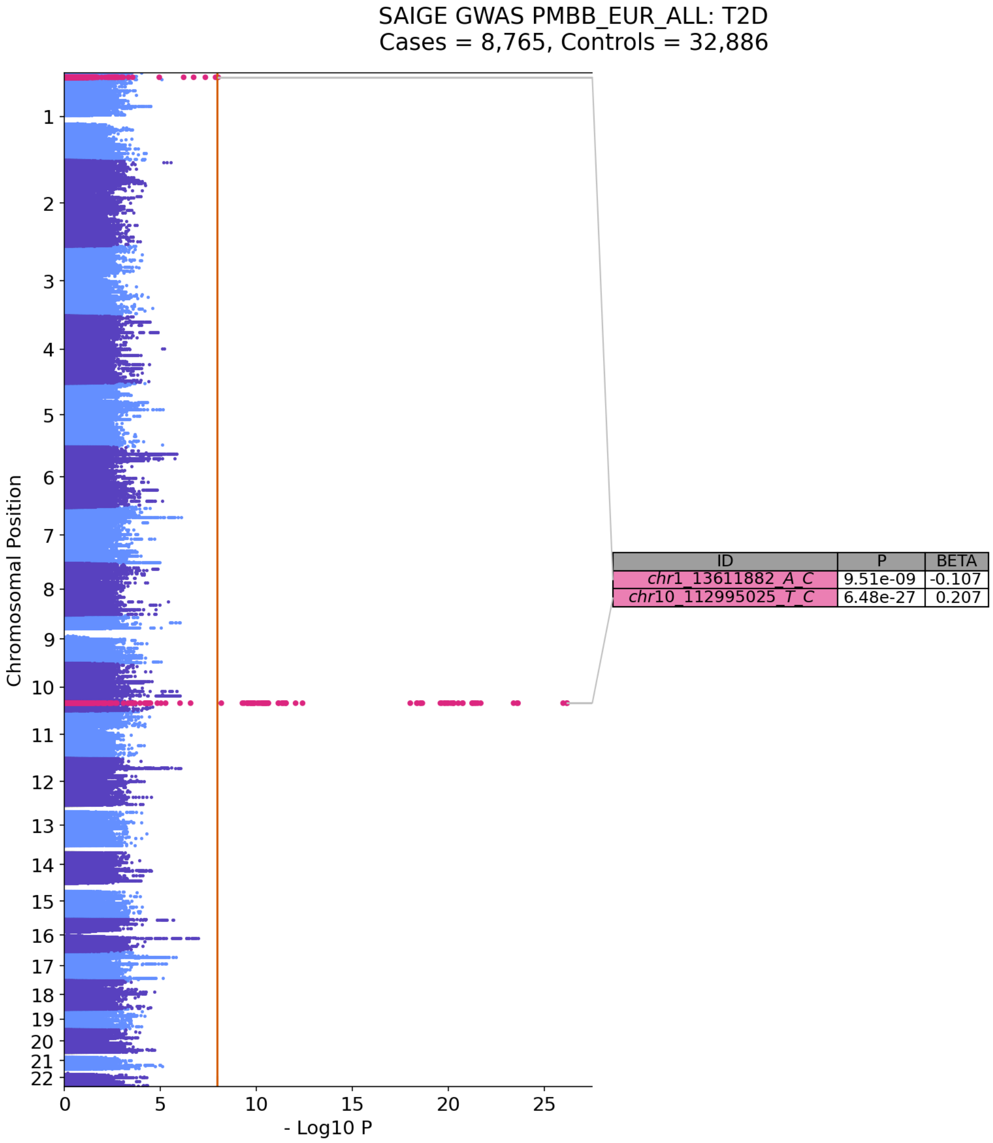


#### Top GWAS Hits

No significant hits found above the p-value threshold.

### GWAS — AAA in PMBB_AFR_ALL

#### GWAS Results — AAA in PMBB_AFR_ALL

##### GWAS Plots — AAA in PMBB_AFR_ALL

Manhattan Plot

QQ Plot

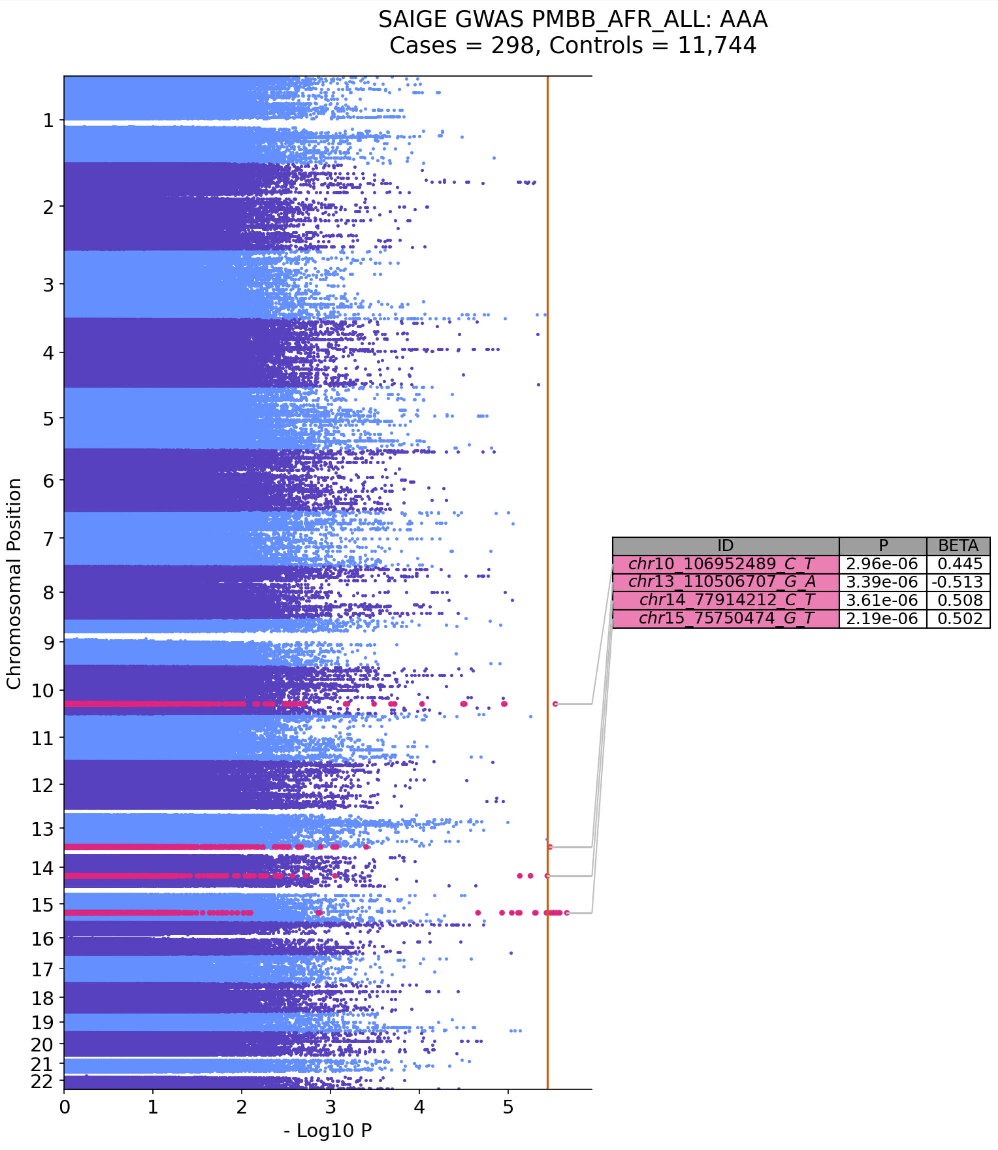


### Top GWAS Hits

No significant hits found above the p-value threshold.

### GWAS — AAA in PMBB_EUR_ALL

#### GWAS Results — AAA in PMBB_EUR_ALL

##### GWAS Plots — AAA in PMBB_EUR_ALL

Manhattan Plot

QQ Plot

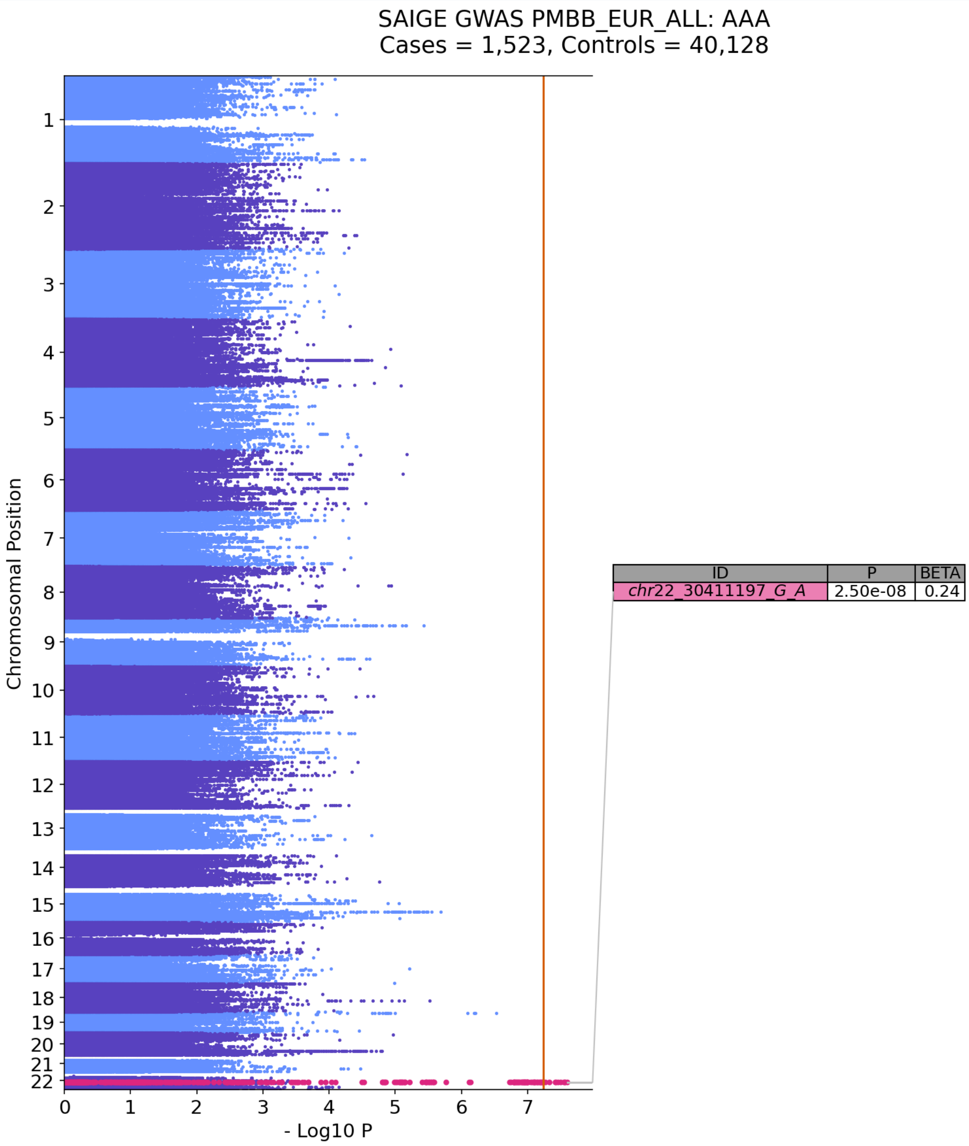


### Top GWAS Hits

No significant hits found above the p-value threshold.

### GWAS — chol_ldl_median in PMBB_AFR_ALL

#### GWAS Results — chol_ldl_median in PMBB_AFR_ALL

##### GWAS Plots — chol_ldl_median in PMBB_AFR_ALL

Manhattan Plot

QQ Plot

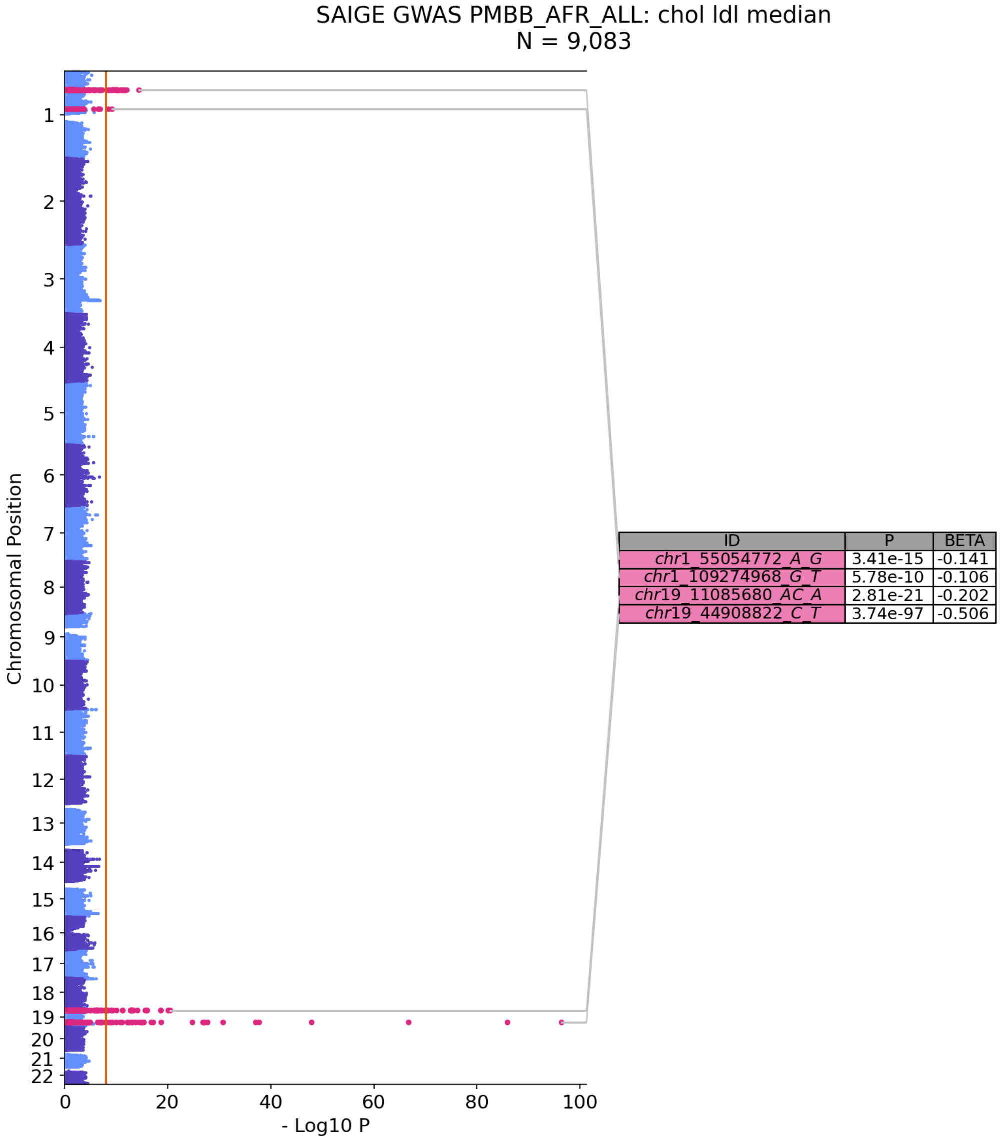


### Top GWAS Hits

No significant hits found above the p-value threshold.

### GWAS — chol_ldl_median in PMBB_EUR_ALL

#### GWAS Results — chol_ldl_median in PMBB_EUR_ALL

##### GWAS Plots — chol_ldl_median in PMBB_EUR_ALL

Manhattan Plot

QQ Plot

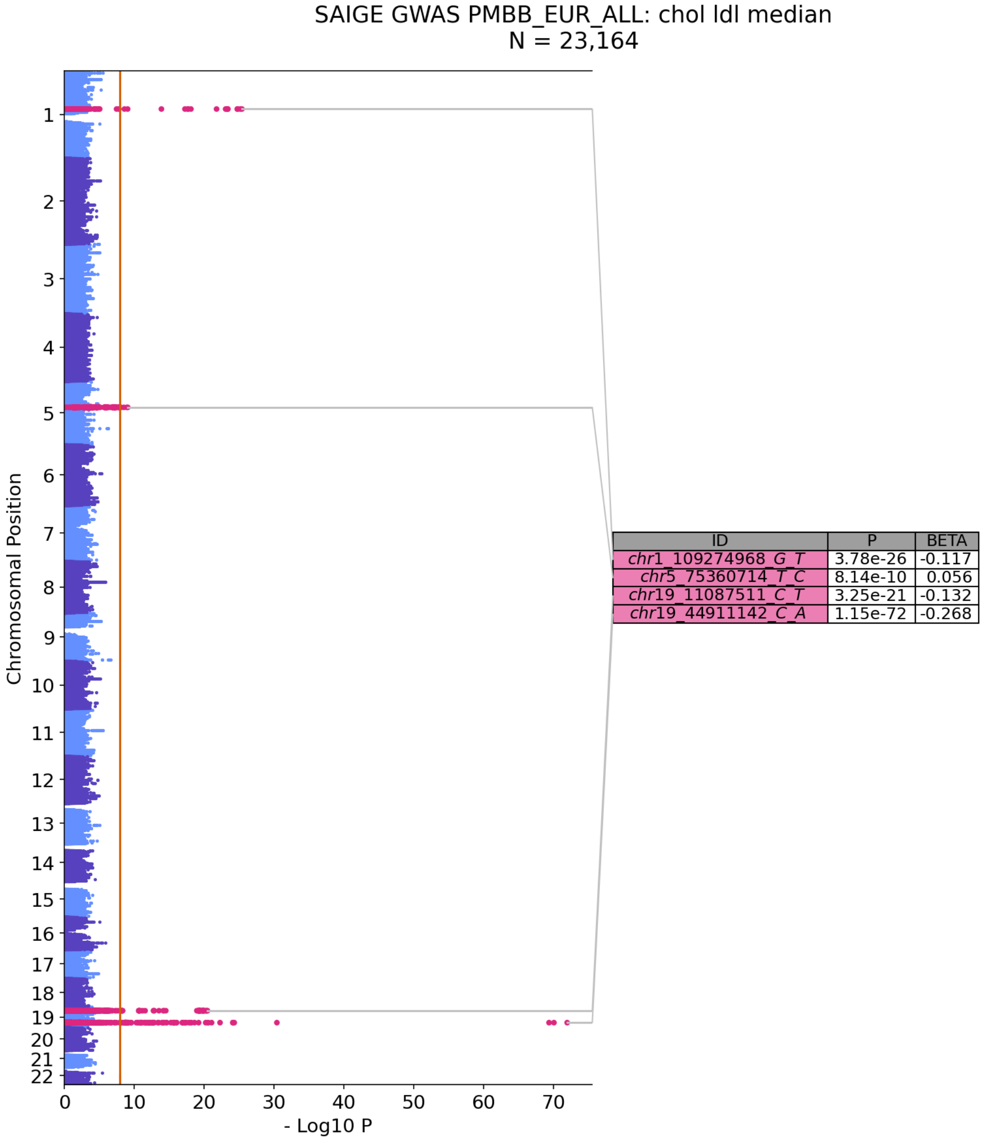


### Top GWAS Hits

No significant hits found above the p-value threshold.

### GWAS — bmi_median in PMBB_AFR_ALL

#### GWAS Results — bmi_median in PMBB_AFR_ALL

##### GWAS Plots — bmi_median in PMBB_AFR_ALL

Manhattan Plot

QQ Plot

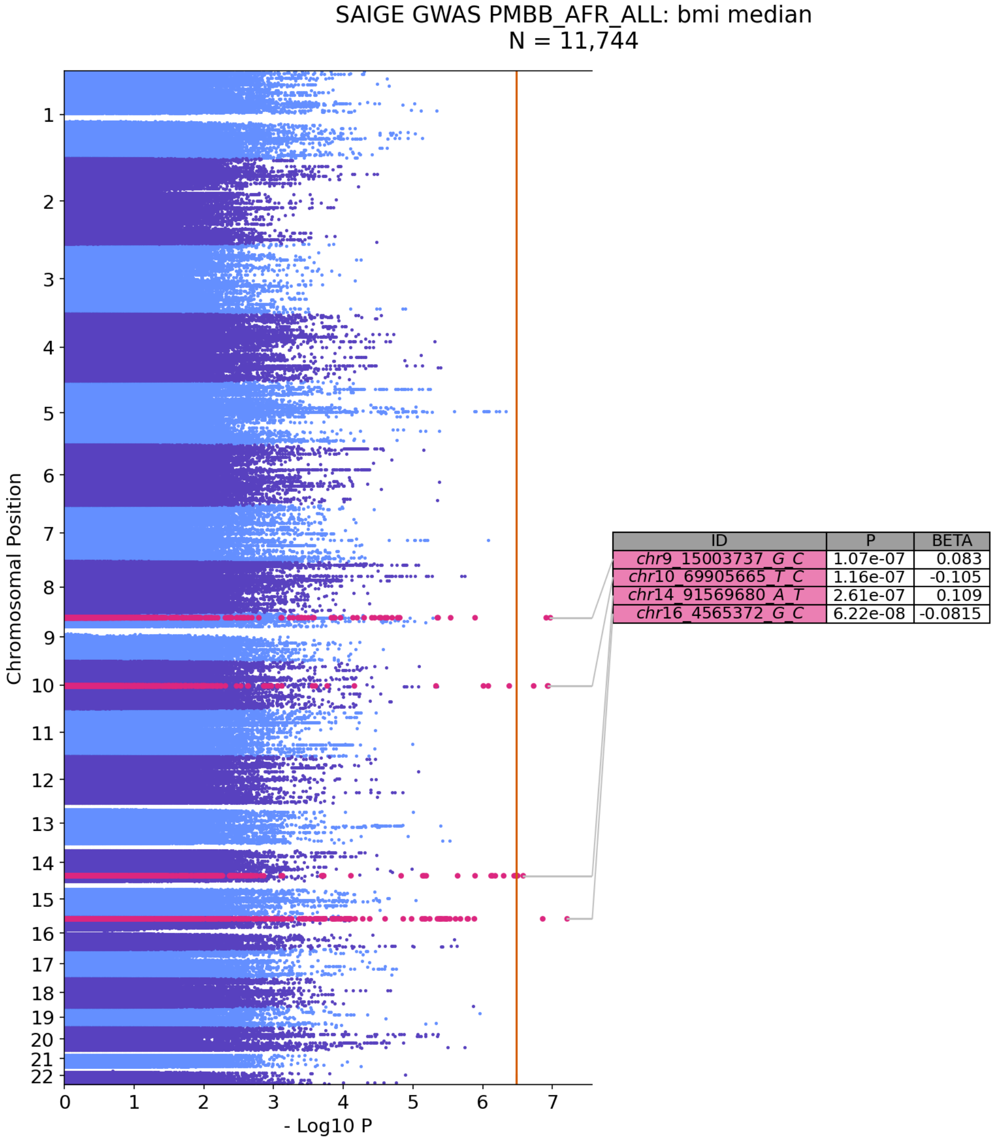


### Top GWAS Hits

No significant hits found above the p-value threshold.

### GWAS — bmi_median in PMBB_EUR_ALL

#### GWAS Results — bmi_median in PMBB_EUR_ALL

##### GWAS Plots — bmi_median in PMBB_EUR_ALL

Manhattan Plot

QQ Plot

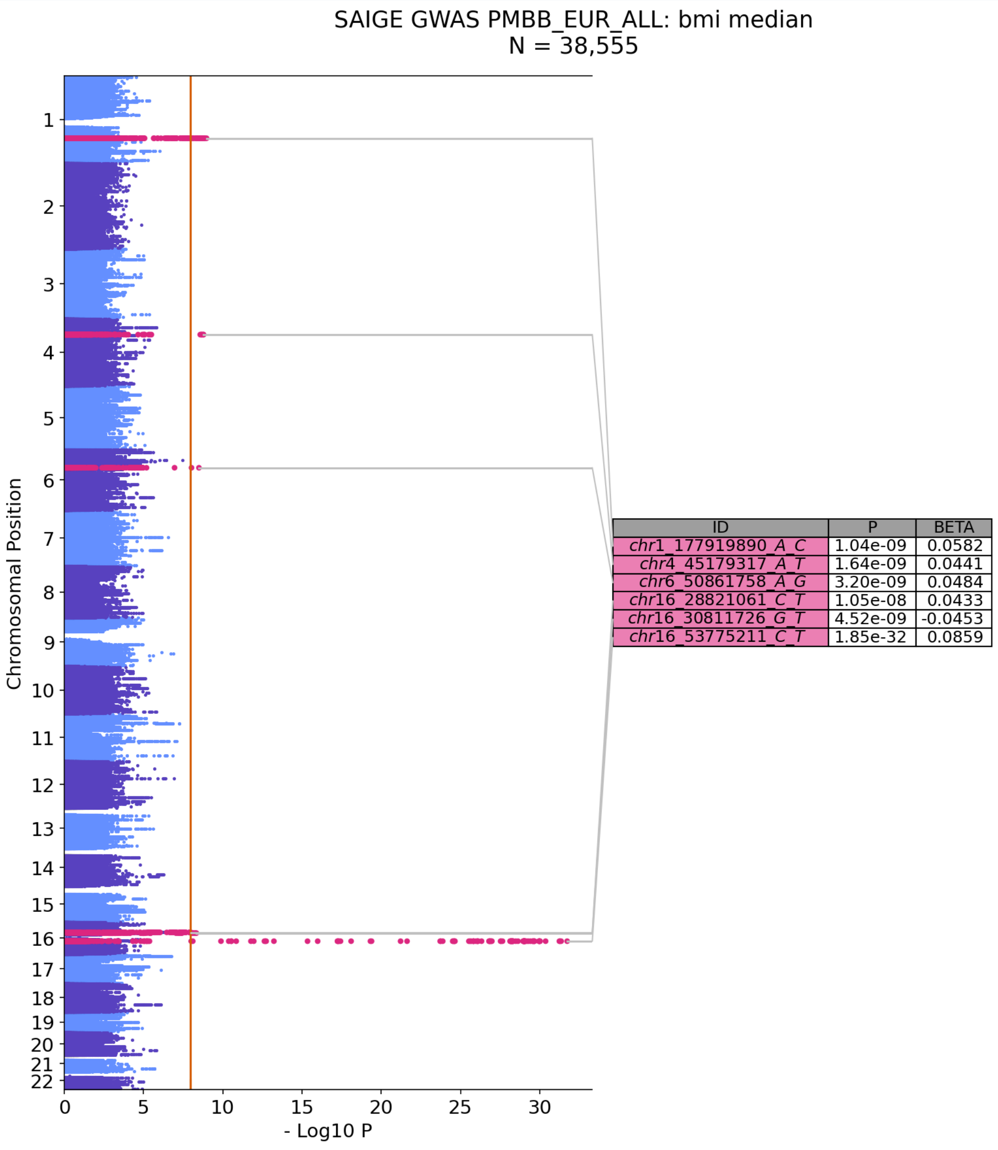


### Top GWAS Hits

No significant hits found above the p-value threshold.

### Analysis Logs

#### SAIGE GWAS Methods Summary

##### Run Info

| Parameter | Value |
|---|---|
| **Run As** | nextflow run /project/pmbb_codeworks/projects/geno_pheno_workbench_dev/SAIGE_FAMILY/workflows/saige_gwas.nf -c ../input/nextflow.conf -c saige_gwas_full.conf -c ../input/special_config.conf -profile cluster -with-report -with-timeline -with-dag -resume |
| **Run Location** | /project/pmbb_codeworks/projects/pmbb-nf-toolkit-benchmarking/nf-saige-gwas-full |
| **Started At** | 2026-03-26T15:42:31.564820174-04:00 |
| **Python Exe** | python |

### Cohorts Phenotypes Chromosomes

| Parameter | Value |
| --- | --- |
| **Cohort List** | PMBB_AFR_ALL, PMBB_EUR_ALL |
| **Sex Strat Cohort List** | — |
| **Bin Pheno List** | T2D, AAA |
| **Quant Pheno List** | chol_ldl_median, bmi_median |
| **Survival Pheno List** | — |
| **Sex Specific Pheno File** | — |
| **Chromosome List** | 1, 2, 3, 4, 5, 6, 7, 8, 9, 10, 11, 12, 13, 14, 15, 16, 17, 18, 19, 20, 21, 22 |
| **Cat Covars** | SEX, Batch |
| **Cont Covars** | AGE, imputed_PC1, imputed_PC2, imputed_PC3, imputed_PC4, imputed_PC5, imputed_PC6, imputed_PC7, imputed_PC8 |
| **Sex Strat Cat Covars** | — |

| Parameter | Value |
| --- | --- |
| **Sex Strat Cont Covars** | AGE, imputed_PC1, imputed_PC2, imputed_PC3, imputed_PC4, imputed_PC5, imputed_PC6, imputed_PC7, imputed_PC8 |
| **Data Csv** | /project/verma_shared/projects/pmbb_release_v3/Imputed/gwas/Input/phenotypes_and_covars_imputed.csv |
| **Cohort Sets** | /project/verma_shared/projects/pmbb_release_v3/Imputed/gwas/Input/cohort_membership_binary.csv |
| **Min Bin Cases** | — |
| **Min Quant N** | — |
| **Min Survival Cases** | 100 |

#### Input File Prefixes

| Parameter | Value |
| --- | --- |
| **Ftype** | BGEN |
| **Use Sparse Grm** | False |

| Parameter | Value |
| --- | --- |
| **Step1 Sparse Grm** | — |
| **Step1 Sparse Grm Samples** | — |
| **Step1 Plink Prefix** | /static/PMBB/PMBB-Release-2024-3.0/Imputed/common_snps_LD_pruned/PMBB-Release-2024-3.0_genetic_imputed.commonsnps |
| **Step2 Plink Prefix** | — |
| **Step2 Bgen Prefix** | /project/verma_shared/projects/pmbb_release_v3/Imputed/chunked/combined/PMBB-Release-2024-3.0_genetic_imputed.chr |
| **Bgen Samplefile** | /static/PMBB/PMBB-Release-2024-3.0/Imputed/common_snps_LD_pruned/PMBB-Release-2024-3.0_genetic_imputed.commonsnps.samplelist.txt |

#### Chunking Parameters

| Parameter | Value |
|---|---|
| **Enable Chunking** | True |
| **Full Chromosome List** | 1, 2, 3, 4, 5, 6, 7, 8, 9, 10, 11, 12, 13, 14, 15, 16, 17, 18, 19, 20, 21, 22 |
| **Chunks List** | /project/verma_shared/projects/pmbb_release_v3/Imputed/gwas/Input/bgen_chunk_list.txt |

#### Saige Step1 Qc

| Parameter | Value |
|---|---|
| **Maf** | 0.01 |
| **Geno** | 0.01 |
| **Hwe** | 1e-06 |
| **Max Vars For Grm** | 200000 |
| **Min Vars For Grm** | 20000 |

| Parameter | Value |
|---|---|
| **Min Rare Vars For Grm** | — |
| **Pruning R2 For Grm** | — |

#### Saige Gwas Parameters

| Parameter | Value |
|---|---|
| **Is Firth Beta** | — |
| **Pcutoffforfirth** | 0.1 |
| **Loco** | TRUE |
| **Gwas Col Names** | CHR → chromosome<br>POS → base_pair_location<br>MarkerID → variant_id<br>Allele1 → other_allele<br>Allele2 → effect_allele<br>AC_Allele2 → effect_allele_count<br>AF_Allele2 → effect_allele_frequency<br>MissingRate → missing_rate<br>BETA → beta<br>SE → standard_error<br>Tstat → t_statistic<br>var → variance<br>p.value → p_value<br>p.value.NA → p_value_na<br>Is.SPA → is_spa_test<br>AF_case → allele_freq_case |

| Parameter | Value |
| --- | --- |
| | AF_ctrl → allele_freq_ctrl<br>N_case → n_case<br>N_ctrl → n_ctrl<br>N_case_hom → n_case_hom<br>N_case_het → n_case_het<br>N_ctrl_hom → n_ctrl_hom<br>N_ctrl_het → n_ctrl_het |
| **Annotate** | False |

#### Survival Parameters

| Parameter | Value |
| --- | --- |
| **Event Time Col** | — |
| **Event Time Bin** | — |

#### Other Parameters

| Parameter | Value |
| --- | --- |
| **Host** | LPC |

| Parameter | Value |
|---|---|
| Gpu | OFF |
| Step1 Script | /usr/local/bin/step1_fitNULLGLMM.R |
| Step2 Script | /usr/local/bin/step2_SPAtests.R |
| Pheno File Id Col | PMBB_ID |
| P Cutoff Summarize | 1e-05 |
| My Bgenix | /opt/conda/bin/bgenix |

Nextflow Report

### Nextflow workflow report

#### [backstabbing_hugle] *(resumed run)*

Workflow execution completed successfully!

##### Run times

26-Mar-2026 15:42:31 - 26-Mar-2026 15:53:07 (duration: **2h 10m 35s**)

... 7459 cached

##### Nextflow command

```
nextflow run
/project/pmbb_codeworks/projects/geno_pheno_workbench_dev/SAIGE_FAMILY/
-c ../input/nextflow.conf -c saige_gwas_full.conf -c
../input/special_config.conf -profile cluster -with-report -with-
timeline -with-dag -resume
```

| | |
|---|---|
| **CPU-Hours** | 294.7 (99.6% cached) |
| **Launch directory** | /project/pmbb_codeworks/projects/pmbb-nf-toolkit-benchmarking/nf-saige-gwas-full |
| **Work directory** | /project/pmbb_codeworks/projects/pmbb-nf-toolkit-benchmarking/nf-saige-gwas-full/work |
| **Project directory** | /project/pmbb_codeworks/projects/geno_pheno_workbench_dev/SAIGE_FAMILY/workflows |
| **Script name** | saige_gwas.nf |
| **Script ID** | 60e0ca240d2e03d479fcd7ffdfa2ad64 |
| **Workflow session** | 8704f5d2-9f45-4f67-bf22-db271653dc8a |
| **Workflow profile** | cluster |
| **Workflow container** | /project/pmbb_codeworks/tools/geno_pheno_toolkit/pmbb-nf-toolkit-saige-family/saige.sif |
| **Container engine** | singularity |
| **Nextflow version** | version 24.04.3, build 5916 (09-07-2024 19:35 UTC) |

Nextflow Report

### Resource Usage

These plots give an overview of the distribution of resource usage for each process.

#### CPU

Raw Usage | % Allocated

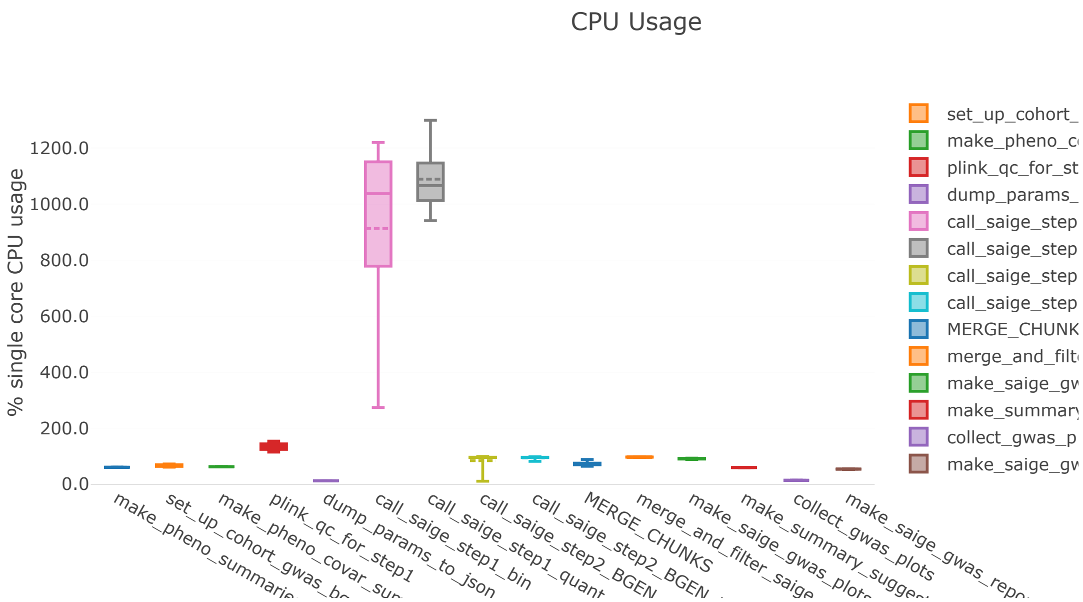


#### Memory

Physical (RAM) | Virtual (RAM + Disk swap) | % RAM Allocated

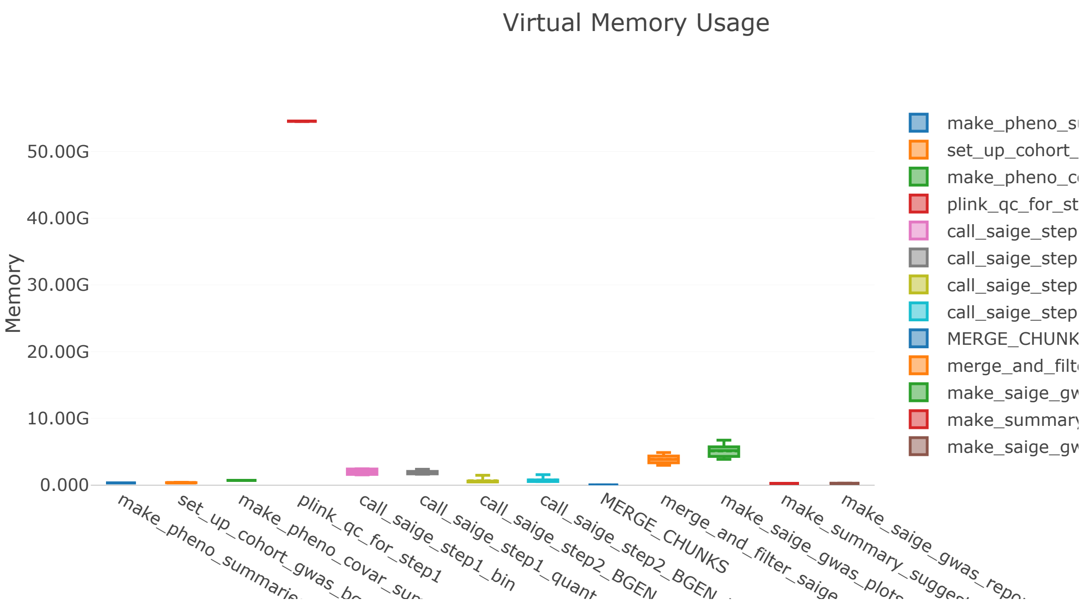


#### Job Duration

Raw Usage | % Allocated

Nextflow Report

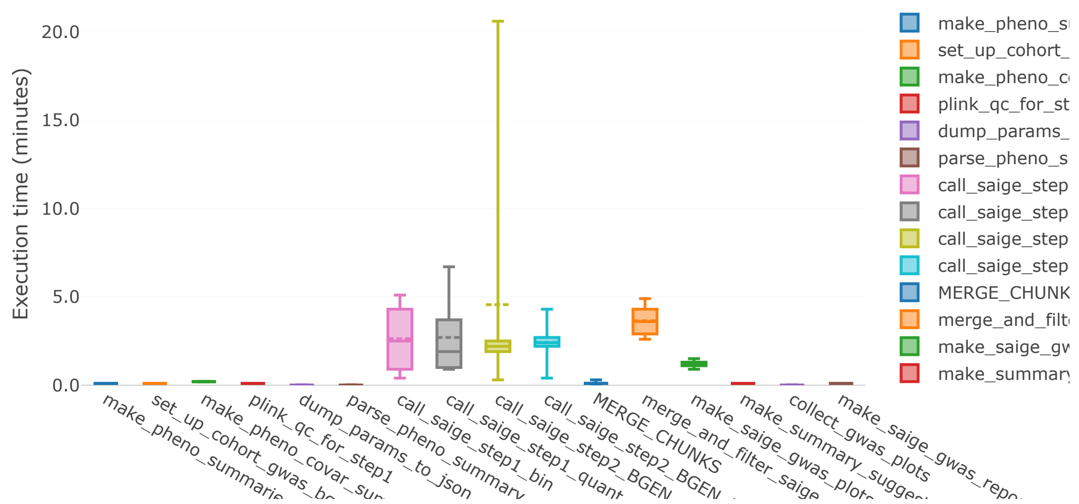


## I/O

Read Write

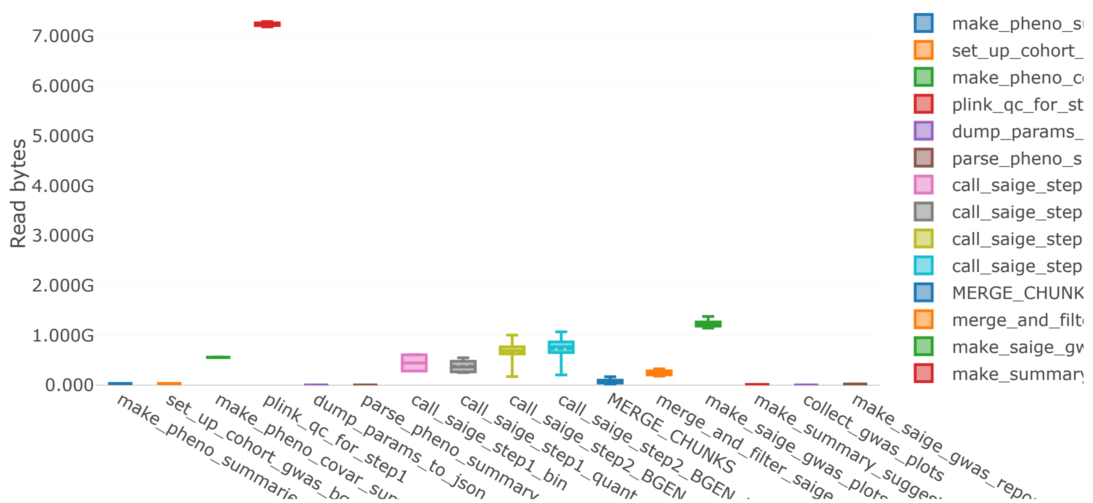


### Tasks

This table shows information about each task in the workflow. Use the search box on the right to filter rows for specific values. Clicking headers will sort the table by that value and scrolling side to side will reveal more columns.

Values shown as: Human readable

Show 25 entries

Filter: Metrics Metadata All Search:

Nextflow Report

| | | | |
|---|---|---|---|
| 1 | SAIGE_GWAS:SAIGE_PREPROCESSING | - | CACHED |
| 2 | SAIGE_GWAS:SAIGE_PREPROCESSING | - | CACHED |
| 3 | SAIGE_GWAS:SAIGE_PREPROCESSING | - | CACHED |
| 4 | SAIGE_GWAS:dump_params_to_json | - | COMPLETED |
| 5 | SAIGE_GWAS:SAIGE_PREPROCESSING | - | CACHED |
| 6 | SAIGE_GWAS:SAIGE_PREPROCESSING | - | COMPLETED |
| 7 | SAIGE_GWAS:SAIGE_STEP1:plink_qc_fc | - | CACHED |
| 8 | SAIGE_GWAS:SAIGE_STEP1:plink_qc_fc | - | CACHED |
| 9 | SAIGE_GWAS:SAIGE_STEP1:call_saige_ | - | CACHED |
| 10 | SAIGE_GWAS:SAIGE_STEP1:call_saige_ | - | CACHED |
| 11 | SAIGE_GWAS:SAIGE_STEP1:call_saige_ | - | CACHED |
| 12 | SAIGE_GWAS:SAIGE_STEP1:call_saige_ | - | CACHED |
| 13 | SAIGE_GWAS:SAIGE_STEP1:call_saige_ | - | CACHED |
| 14 | SAIGE_GWAS:SAIGE_STEP1:call_saige_ | - | CACHED |
| 15 | SAIGE_GWAS:SAIGE_STEP1:call_saige_ | - | CACHED |
| 16 | SAIGE_GWAS:SAIGE_STEP1:call_saige_ | - | CACHED |
| 17 | SAIGE_GWAS:SAIGE_VAR_STEP2:call_sa | - | CACHED |
| 18 | SAIGE_GWAS:SAIGE_VAR_STEP2:call_sa | - | CACHED |
| 19 | SAIGE_GWAS:SAIGE_VAR_STEP2:call_sa | - | CACHED |
| 20 | SAIGE_GWAS:SAIGE_VAR_STEP2:call_sa | - | CACHED |

Nextflow Report

| | | | |
|---|---|---|---|
| 22 | SAIGE_GWAS:SAIGE_VAR_STEP2:call_sa | - | CACHED |
| 23 | SAIGE_GWAS:SAIGE_VAR_STEP2:call_sa | - | CACHED |
| 24 | SAIGE_GWAS:SAIGE_VAR_STEP2:call_sa | - | CACHED |
| 25 | SAIGE_GWAS:SAIGE_VAR_STEP2:call_sa | - | CACHED |

Showing 1 to 25 of 7,659 entries

Previous 1 2 3 4 5 … 307 Next

Generated by Nextflow, version 24.04.3
